# The induction of systemic resistance to barley powdery mildew by rhizosphere bacterial communities does not disrupt the structure or function of native microbial communities

**DOI:** 10.64898/2026.01.21.700805

**Authors:** Linda Rigerte, Anna Sommer, A. Corina Vlot, Luis Daniel Prada-Salcedo, Thomas Reitz, Anna Heintz-Buschart, Mika T. Tarkka

## Abstract

Synthetic microbial communities (SynComs) could help plants withstand biotic stress and reduce the need for pesticides. With this in mind, we created two SynComs, comprising bacterial strains isolated from the rhizospheres of barley and wheat. We then studied their potential to trigger induced systemic resistance against the barley pathogen *Blumeria graminis* f. sp. *hordei (Bgh)*. To investigate the plant-microbial interactions from the perspective of both plants and microbes, we performed DAF staining to quantify *Bgh* propagation in plant leaves, analysed leaf transcriptomes and conducted rhizosp here 16S rRNA gene metabarcoding and rhizosphere metatranscriptome analysis. Our results demonstrate that the SynComs elicit defence responses in barley against *Bgh* in a manner similar to that of the positive control strain *Pseudomonas simiae* WCS417r. The SynComs act without triggering a strong gene response prior to inoculation with the plant pathogen or affecting plant-associated prokaryote communities; they only mildly influence bacterial gene expression in the rhizosphere. Instead, they act as priming agents, preparing the plant for further pathogen attack. These findings suggest that protective SynComs can be applied in the field without causing signficant disruption to native microbial communities.

## 2. Introduction

Plant-associated microbial communities have a positive effect on a plant’s fitness, development, and tolerance of stress (Mendes et al. 2013; Panke-Buisse et al. 2015; Lau and Lennon 2012). Plant beneficial bacteria (PBB) have been isolated and characterized from the root, leaf and stem surfaces, and inside plant tissues (Glick 2014). They mobilize nutrients (Israr Asghar 2023), produce phytohormones such as indole-3-acetic acid, ethylene and jasmonic acid (Kejela 2023) and improve plant immunity and resistance against pathogens (Fujiwara et al. 2016). Generating host-associated microbial culture collections has enabled plant growth promoting (PGP) bacteria to be used as bioinoculants in the form of single strains or as consortia via the synthetic microbial community (SynCom) approach (Niu et al. 2017). SynComs are constructed microbial consortia, usually composed of selected commensal strains with PGP abilities. They can be used to rebuild a host microbiome under controlled conditions and make it possible to study the interactions between microbes and their hosts, as well as interactions inside the microbial community (Northen et al. 2024).

Plants possess a complex innate immune system to defend against pathogenic bacteria, fungi, protists, nematodes and viruses (Agrios 2009). When attacked by a pathogen, plant systemic signaling transfers the information from the affected parts through the plant to tissues that are still unaffected and thus triger a coordinated response (Vlot et al. 2021). More precisely, plant cells use pattern recognition receptors to recognize the microbe or pathogen–associated molecular patterns. This process provokes pattern-triggered immunity, which together with effector-trigerred immunity can induce salicylic acid signalling and activate systemic acquired resistance (SAR) (Vlot et al. 2021; Pieterse et al. 2012).

Beneficial bacteria can mediate induced systemic resistance (ISR), a systemic immunity pathway distinct from SAR, by priming the aforementioned defence mechanisms, a phenomenon first observed with *Pseudomonas* and *Bacillus* strains (van Wees et al. 2008). This priming puts the plants under an “alerted” state, enabling them to respond more quickly and/or strongly to subsequent pathogen infections (Heil 2001; Conrath et al. 2015; Martinez-Medina et al. 2016). Priming is suggested to be effective against a wide range of pathogens, including (hemi-)biotrophic and necrotrophic types, and even herbivores (Vlot et al. 2021). ISR by commensal *Pseudomonas* strains has been shown to depend on jasmonic acid and ethylene signaling pathways (Vlot et al. 2021; Pieterse et al. 2012; Yu et al. 2022), whereas the *Bacillus-*and *Streptomyces*-elicited pathways may also employ salicylic acid signalling (Kloepper et al. 2004; Ebrahimi-Zarandi et al. 2022; van Wees et al. 2008; Kurth et al. 2014). ISR results in either subtle (Heil and Bostock 2002) or extensive (Kurth et al. 2014) changes in gene expression in the unaffected tissues that primes the plant for an improved defense response upon pathogen attack. Despite ISR mainly elicited in the roots through interaction with beneficial rhizobacteria, it can protect above-ground tissues (like leaves) by priming JA- and ethylene-dependent defence pathways (Pieterse et al. 1998; van Wees et al. 2008). Currently, it is unclear which bacterial taxa or communities are capable of inducing ISR in barley, and whether such responses are specific to particular biotic or abiotic stress conditions.

*Blumeria graminis* f. sp. *hordei (Bgh)* is a widespread fungal pathogen that causes the fungal disease powdery mildew. This affects a wide range of plants including cereal crops like barley (Sacharow et al. 2023) and is reported to occur in several barley growing regions (Wang et al. 2023; Cieplak et al. 2022; Cowger et al. 2018; Kloppe et al. 2022). Powdery mildew affects plant leaves and can adversely impact crop yield (Czembor and Czembor 2023) and photosynthesis (Coghlan and Walters 1992), particularly under moist and warm weather conditions which favor the pathogen (Elagamey et al. 2023; Bhardwaj et al. 2021). Efforts to study ISR have often employed using the bacterial strain *Pseudomonas simiae* WCS417r, which was recently shown to protect barley from Bgh (Pieterse et al. 2021, Sommer et al. 2026). This robust rhizobacterium has been shown to confer ISR in various host plants (Pieterse et al. 2021; Zhu et al. 2022) and is used as a positive control in this study.

The negative impacts of climate change, such as rising temperatures and prolonged droughts, strongly affect plants (Sato et al. 2024). Prolonged drought, for instance, make plants more susceptible to disease by reducing photosynthesis, transpiration and nutrient uptake (Hatfield and Prueger 2015; Nievola et al. 2017). Furthermore, climate change may support plant disease and pest dispersal (Delgado-Baquerizo et al. 2020), thus making plants more susceptible and cause damage and reduce yields (Rezaei et al. 2023; Singh et al. 2023). Such challenges suggest the need for sustainable approaches that enhance plant resilience to abiotic and biotic stress, thus ISR can work as an important tool to maintain plant health, agricultural output, and environmental sustainability under future climatic conditions.

ISR in plants has been widely described in model species like *A. thaliana* (Weller et al. 2012; Ilham et al. 2019; van Wees et al. 2000) and tomato (Suresh et al. 2022; Hu et al. 2018; Mehari et al. 2015). Even though interactions between barley and *Bgh* have been extensively studied, including host–pathogen signaling and defense responses, comparatively few studies have addressed ISR in this pathosystem, particularly in the context of defined beneficial microbial communities. While recent work has started to investigate ISR triggered by barley-native microbial communities, comparative studies using well-defined synthetic communities from different host species (e.g., barley vs. wheat) remain limited. Bziuk et al. (2022) demonstrated that a suspension of barley rhizosphere microbiome can enhance the immune response and reduce the incidence of powdery mildew disease. Building on this, we tested whether SynComs from the rhizospheres of barley and wheat, composed of bacterial strains enriched under drought stress (each comprising Firmicutes, Proteobacteria, and Actinobacteria), could trigger ISR in barley plants against the same pathogen. The ISR-eliciting bacterium *P. simiae* WCS417r (WCS417r) was used as a positive control. We expected that due to the presence of Firmicutes and Actinobacteria (Kloepper et al. 2004; Ebrahimi-Zarandi et al. 2022) i) both SynComs would elicit an ISR to the same extent as WCS417r, but in a jasmonic acid-, ethylene- and salicylic acid-dependent manner. We also (ii) hypothesized that, given that barley was used as the host plant in our experiment, the barley SynCom would elicit a stronger response than the wheat SynCom due to the plant host preference of rhizosphere microbiomes. Our final hypothesis was that, iii) at harvest time, only a few SynCom members would remain active and abundant in the rhizosphere.

## 3. Materials and Methods

### 3.1. Collection of barley and wheat rhizobacteria

Barley and wheat rhizobacteria were sampled from growing cereals at the Global Change Experimental Facility (GCEF) in Bad Lauchstädt (51° 23’ 123 30N, 11° 52’ 49E). GCEF is a field research station site of the Helmholtz Centre for Environmental Research (https://www.ufz.de/index.php?en=42385) that has a Haplic Chernozem soil type (Altermann et al. 2005). The site has a temperate continental climate with a mean annual temperature of 9.7°C and mean annual precipitation of 438 mm (1993-2013) (Schädler et al. 2019). The bacterial strains used in this study were sampled from organic and conventional farming plots that were exposed to an ambient and future climate treatment. The latter comprises besides warming a 10% increase in precipitation in spring (March-May) and autumn (September-November), as well as a rainfall reduction by 20% in summer (June-August). This simulates the climate expected for Bad Lauchstädt in 2080 (Schädler et al. 2019). The strains were isolated from the rhizospheres of the barley cultivar “Antonella” and the wheat cultivar “RGT Reform” in two different stages of plant development, stem elongation stage BBCH 37-39 and grain filling stage, BBCH 75-77. The isolation process is described in Breitkreuz et al. (2020) and Rigerte et al. (2025). Briefly, the rhizosphere was collected by gently detaching the soil from the plant roots, after which the organic material was removed manually and by sieving (2 mm). The soil was then mixed and suspended to double-distilled water. The bacteria were separated from the soil by sonication, and a suspension was inoculated on Pikovskaya’s agar medium (Pikovskaya 1948). Plates were then incubated for three weeks at 25°C and colonies showing phosphate solubilization (observed as clear halos around them) were purified.

### 3.2. Selection of bacterial isolates

Polyethylene glycol 8000 (PEG) was used to lower the water potential of agar medium and thereby assess the drought tolerance of the isolated strains. Drought stress was simulated by applying a 500 g/L solution of PEG (PEG0.5), which corresponds to an osmotic potential of approximately −1.1 MPa (Verslues et al. 2006). Drought tolerance was assessed by calculating the percentage reduction in colony diameter under PEG0.5 compared to PEG0. Genomic DNA was extracted by adding PEG to the bacterial suspensions and then vortexed with glass beads according to the method of Breitkreuz et al. (2019). DNA amplification was performed using PCR with the primers 27F and 1492R (Lane 1991) and the Promega Green system (Promega, Madison, WI, USA). Subsequently, partial 16s rRNA gene sequences were obtained by Sanger sequencing using the primer BAC 341F at LGC Genomics (Berlin, Germany). The main objective of Sanger sequencing was to obtain sequence regions overlapping with ASVs obtained by 16s rRNA gene amplicon sequencing in previous studies. These amplicon sequencing darasets were generated from barley and wheat rhizosphere samples and were used as a referene for selecting bacterial strains for SynCom construction. Amplicon sequence variants (ASVs) that were more frequent under future climate conditions compared to ambient conditions in barley and wheat rhizosphere samples (Breitkreuz et al. 2021a), as well as ASVs enriched under drought compared to well-watered conditions in a greenhouse experiment using the same chernozem soil (Breitkreuz et al. 2021b), were selected from these datasets. These ASVs were then compared to the 16S rDNA sequences from our cultured bacterial collection to find isolates with identical or highly similar sequences. From this group, sixteen PEG-tolerant strains that showed similar growth on PEG0 and PEG0.5 were selected to assemble the Drought-Tolerant Synthetic Community (DT-SynCom). A full overview of the study workflow is presented in Figure 1.

**Figure 1.**
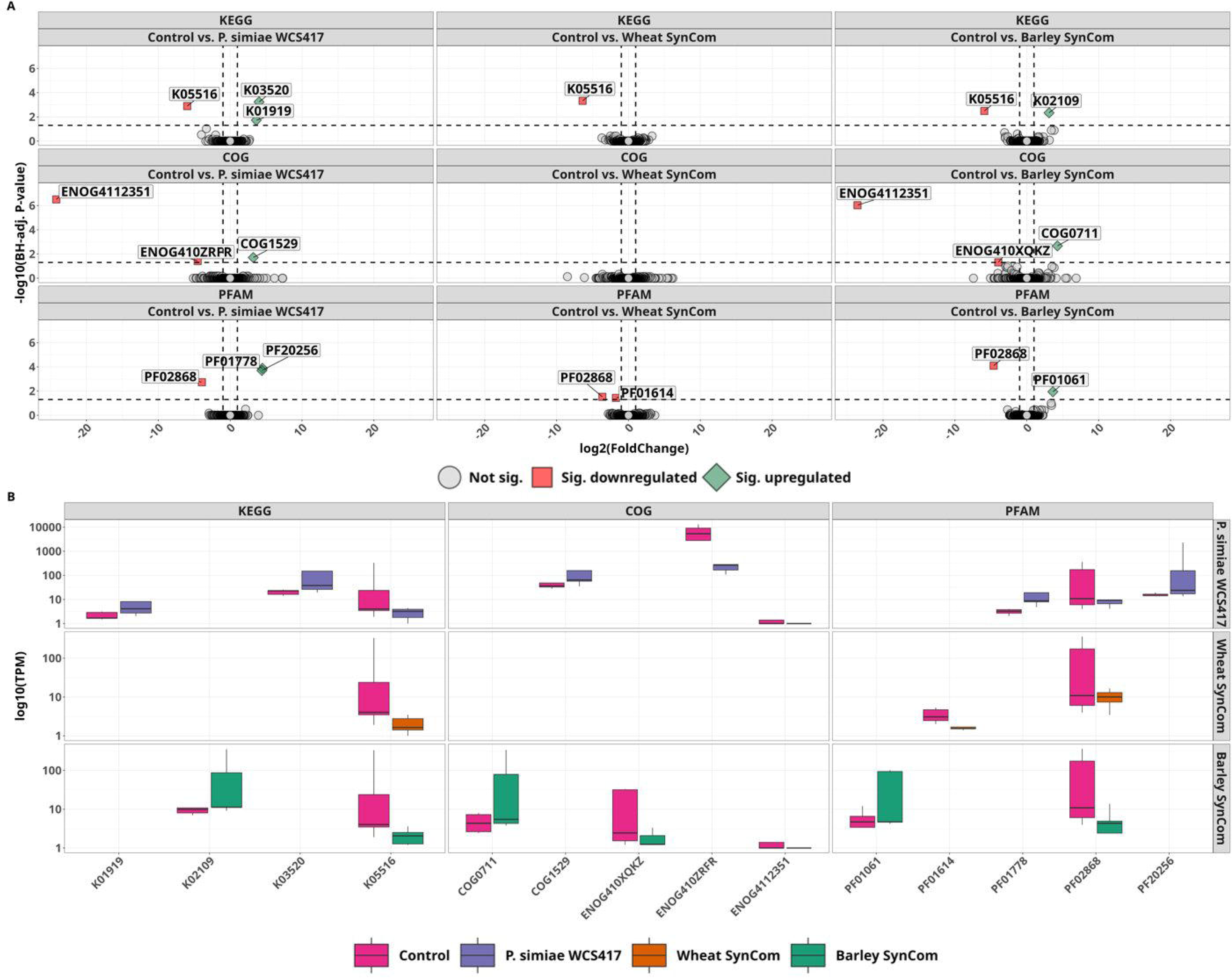
**Workflow of the study**, illustrating the work described in the Materials and methods section 3.1, 3.2 and 3.5 (created in BioRender).

### 3.3 Composition of a synthetic microbial community (SynCom) from barley and wheat

The final synthetic communities consisted of bacterial strains that were isolated from the rhizosphere of barley and wheat. The strains were selected based on their drought tolerance and 16S rDNA sequence identity as described in sections 3.1 – 3.2 The bacterial strains included in both SynComs were described in a previous study (Rigerte et al. 2025) and are listed in Table 1 to document the methods used in this work. Each SynCom was constructed by combining equal volumes of single-strain cultures as described in the following section.

**Table 1.**
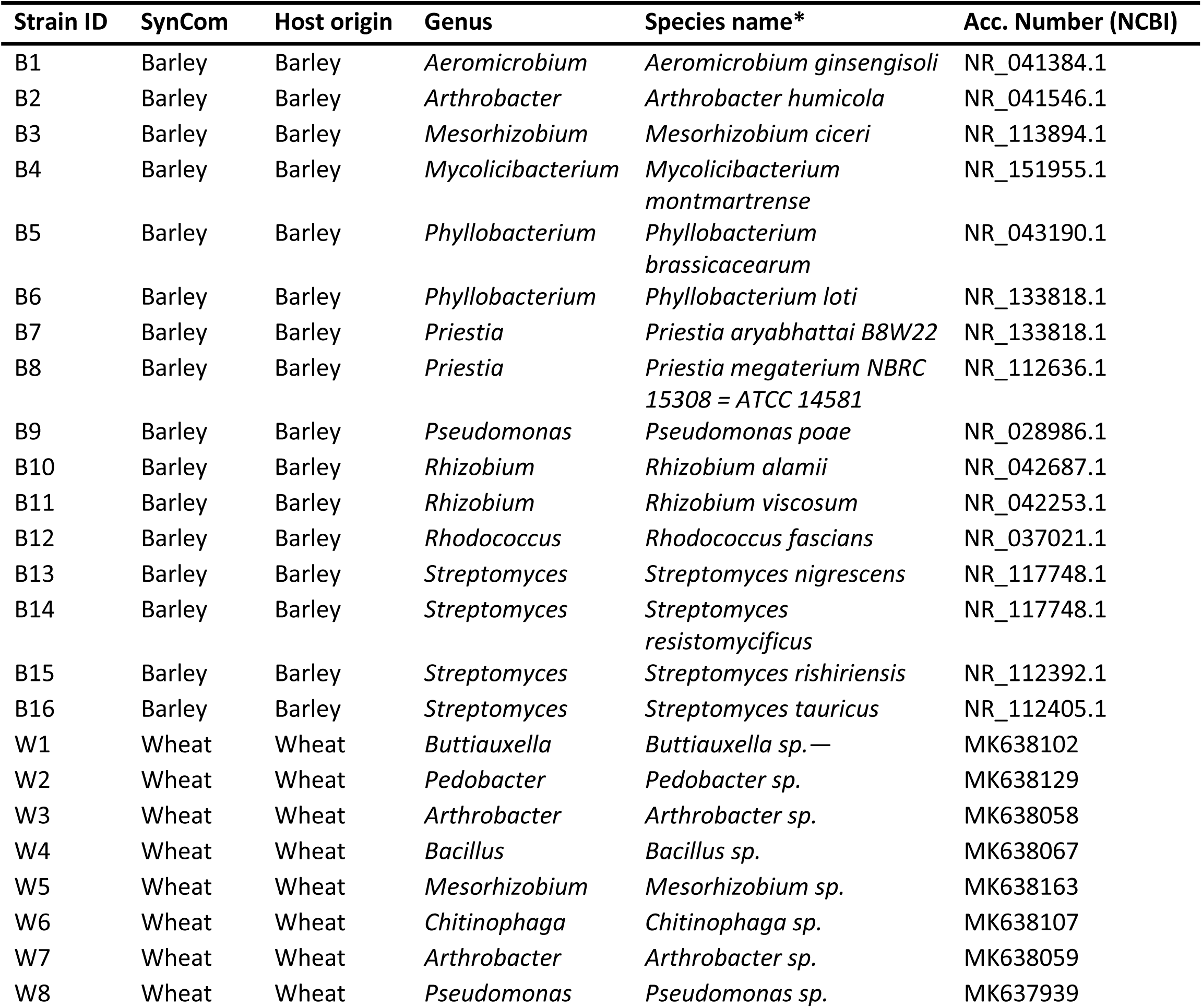
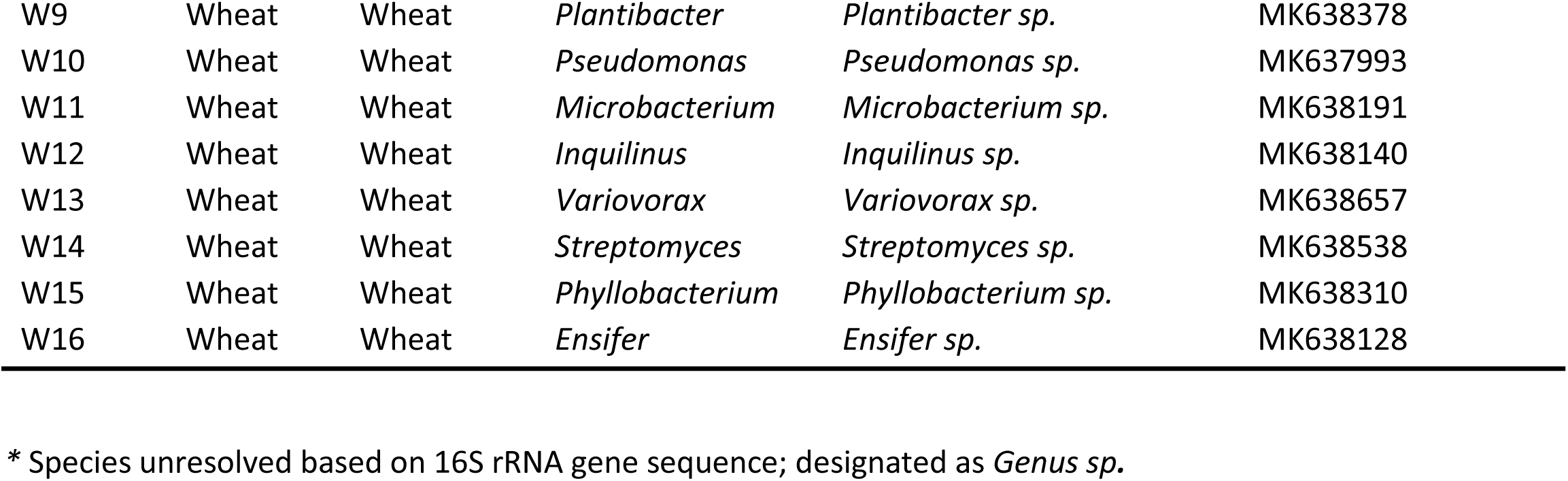
Composition of barley and wheat SynComs (refer Materials and Methods for details.

### 3.4. Preparation of the drought-tolerant synthetic community (DT-SynCom)

Bacterial strains from the rhizosphere of barley and wheat were cultivated for three days at 25 °C on yeast malt extract (YME) agar. The YME medium was prepared by dissolving the constituent ingredients (7 g malt extract, 2.8 g yeast extract, 10.5 g agar (2%) and 2.8 g glucose) in 700 ml water and autoclaving for 20 minutes. Each strain was resuspended in 10 mM magnesium chloride (MgCl_2_) until the optical density (OD) at 600 nm reached a value of 0.5. All individual suspensions were then centrifuged, the supernatant decanted and the bacterial pellets resuspended in 1 ml MgCl_2_. The barley strains were combined in equal proportions into a synthetic community and the wheat strains into a separate community, with each strain adjusted to a final concentration of 10^9^ cfu/ml in 10 mM MgCl_2_.

### 3.5. Propagation of powdery mildew

Barley seeds were sterilized (excluding the surface sterilization step with 75% EtOH), sown and grown for three weeks as previously described. Barley plants were then inoculated with *Bgh* as describedbelow (Lenk et al. 2018; Lenk et al. 2019). Plants were then grown for an additional ten days to obtain mature powdery mildew spores, which were used for infections. Propagation was repeated once a week to maintain a viable powdery mildew culture.

### 3.6. Inoculation of barley plants with SynComs and powdery mildew infection assay

Plant inoculation and powdery mildew infection assays were performed following established protocols (Lenk et al. 2018; Lenk et al. 2019), with minor modifications as described below. Barley seeds (variety “Golden Promise”) were first surface sterilized by a one-minute wash with 75% ethanol (25 inversions per minute), followed by a five-minute incubation in sodium hypochlorite (NaClO). The washing step was followed by a ten-minute incubation in sterile water and repeated three times. The seeds were then placed on sterile NB plates (Table S1) and kept in the dark at room temperature for germination. After four days, the seeds were divided into four groups and inoculated by means of a two hours long incubation with one of the following treatments: the wheat SynCom, the barley SynCom, or WCS417r (positive control). Plants were inoculated with either freshly prepared SynComs or previously prepared SynComs that were stored at -80°C in 25% glycerol and washed three times with MgCl_2_ (10 mM) to remove glycerol prior to use. The forth group was inoculated with MgCl_2_ (10 mM) to serve as a mock control solution. The barley plants were then grown for 3 weeks under a 14/10-hour day/night cycle at 20°C/16°C respectively.

On day 21, the rhizosphere of each plant was collected by carefully removing the root-attached soil by hand, and the second and third leaves of three separate plants, were sampled and stored frozen until subsequent nucleic acid extractions. The remaining barley plants and a glass slide were then placed in a cardboard box and infected with *Bgh* by gently shaking the infected plant over the box. To spread the spores, a styrofoam board was fanned ten times in all directions. After one hour, the spores deposited on the glass slide were counted to verify the spore density of approximately 30 spores per mm^2^. One week later, the plant leaves were harvested to evaluate the infection intensity.

### 3.7. Plant diaminofluorescein (DAF) staining

Plant diaminofluorescein staining was performed following established protocols (Lenk et al. 2018; Lenk et al. 2019). The MES-KOH (morpholinoethanesulfonic acid-potassium hydroxide) 0.5 M stock solution was prepared by adding 21.32 g MES to 200 ml H_2_O and adjusting the pH to 5.7 with KOH. The bottle was then wrapped in an aluminum foil, as MES is light-sensitive, and then placed in the refrigerator till further use (shelf life of the solution is one month). The prepared MES stock solution was then used to prepare the working solution DAF-FM-DA described in Table S2. For staining, the second and third leaf of barley were selected, and 16 leaf discs per treatment of inoculated plants were immersed in 4 ml of working solution (without added DAF) for 45 minutes (in the dark), then the solution was removed and 4 ml of DAF solution was added. The leaf discs were then incubated in the dark for 45 minutes and then vacuum infiltrated 3 to 5 times to aspirate the solution into the plant tissue, followed by incubation under light for 105 minutes. The solution was then removed and 1 ml of working solution was added for a wash step to remove any remaining DAF-FM DA. The leaf discs were then dried on a paper towel and placed upside down on a 96-multiwell plate filled with 1% phytoagar (any meniscus formed by the agar medium was removed by scraping to obtain a uniform surface for fluorescence measurement). The plate was then sealed with qPCR-foil and the Keyence BZ-X800 fluorescence microscope was used to image the leaf discs. Fluorescence was measured using a BZ-X Filter GFP (model OP-87763) with an excitation filter of 470/40 nm, an emission filter of 525/50 nm and a dichroic mirror with a cutoff wavelength of 495 nm. Fluorescence was normalized for each replicate as the ratio of the measured intensity to the mean intensity of the empty wells. A small pseudocount was added for log10 transformation to avoid undefined values when intensity ratios were zero (pseudocount = 0.1 × smallest non-zero ratio observed). Values were then plotted as log10(ratio + pseudocount).

### 3.8. Nucleic acid extraction and sequencing

#### Rhizosphere metatranscriptome

total RNA from barley rhizosphere samples was extracted using the RNeasy PowerSoil Total RNA Kit (Qiagen, Hilden, Germany) following the extraction protocol provided by the manufacturer. Sequencing was performed by Azenta Life Sciences (Leipzig, Germany) on an Illumina instrument after rRNA depletion in a paired-end configuration generating approximately 20 million paired-end reads of 150 bp read length per sample.

#### Host leaf gene expression

Second leaf of barley plant was sampled and grinded using liquid nitrogen. Leaf RNA was extracted using the NucleoSpin RNA Plant Kit (Macherey-Nagel, Düren, Germany) and Agilent Bioanalyzer was used to determine the RNA integrity number (RIN) which was ≥ 7 for all samples. Samples were sequenced by Novogene (Düsseldorf, Germany) and Illumina sequencing was performed using a NovaSeq X Plus Series sequencing platform, which generated approximately 20 million paired-end reads of 150 bp length per sample (40 million reads in total).

#### Rhizosphere 16S rRNA gene metagenomes

barley rhizosphere DNA was extracted using the DNeasy PowerSoil Pro Kit (Qiagen, Hilden, Germany) and barley leaf microbiome DNA was extracted using NucleoSpin Plant II kit (Macherey-Nagel, Düren, Germany). The V4 region of the 16S rRNA gene was amplified using the primers 515F (GTGYCAGCMGCCGCGGTAA) and 806R (GGACTACNVGGGTWTCTAAT) (Caporaso et al. 2011), each with Illumina overhang adapters (P5: TCGTCGGCAGCGTCAGATGTGTATAAGAGACAG; P7: GTCTCGTGGGCTCGGAGATGTGTATAAGAGACAG). The Illumina MiSeq platform with Reagent Kit v3 was used to generate an average of ∼200,000 read pairs per sample (2x 300 bp).

All sequencing data generated in this study, including rhizosphere metatranscriptomes, rhizosphere 16S rRNA gene amplicon sequencing data, and host leaf RNA sequencing data, have been deposited in NCBI under BioProject ID PRJNA1405459.

### 3.9. Rhizosphere 16S rDNA amplicon sequencing bioinformatics and microbiome analysis

The sequenced 150 bp paired-end reads were processed using the Dadasnake pipeline v0.11 (Weißbecker et al. 2020; Callahan et al. 2016) with default parameters. The process included quality filtering and trimming by truncating forward and reverse reads to 170 bp and 130 bp, respectively. Reads with a minimum quality score of 13 were retained and a maximum expected error rate of 0.2 for both read directions was allowed. Error models were learned and sequencing errors corrected using DADA2, which was followed by dereplication, which collapsed identical reads into amplicon sequence variants (ASVs). ASVs were subsequently clustered at 97% sequence identity to generate operational taxonomic units (OTUs) (Schloss 2021; Estensmo et al. 2021). Chimeras were removed using DADA2’s consensus algorithm. Taxonomic classification of the produced OTUs was performed against the SILVA v138.1 database (Quast et al. 2013). Additionally, OTUs that could have been potentially associated with the SynCom strains were identified using MMseqs2’s easy-search module by demanding a minimum sequence identity of 99% and 99% sequence coverage of the OTU sequence by the corresponding 16S rDNA from the SynCom genomes published by our group (Rigerte et al. 2025). OTUs identified as belonging to the following taxa were removed in R v4.4.1 prior to further analysis: chloroplasts, mitochondria, Rickettsiales (obligate intracellular symbionts/parasites), Diploririckettsiales (not known to be plant-associated), Piscirickettsiales (not known to be associated with soils/plants), fungi, algae, Escherichia spp., and taxonomic assignments that were explicitly marked as “unknown” and/or “unclassified”.

Further processing was performed using functions from the phyloseq v1.50 (McMurdie and Holmes 2013), microViz v0.12.7 (Barnett et al. 2021), and vegan v2.7-1 (Oksanen et al. 2025) R packages. The OTU table was pre-processed using the tax_fix() function from microViz to fill in missing taxonomic ranks at higher levels from lower levels. OTUs with zero reads across all samples were removed. Alpha diversity metrics (Shannon, Chao1 and observed) were calculated using the estimate_richness() function from phyloseq without additional filtering of the OTU table. To calculate beta diversity, the data were first filtered to remove OTUs with fewer than five reads across all samples, with no phyla being removed. Samples with zero reads across all OTUs were then filtered out. The data were then rarefied to even depth using phyloseq’s rarefy_even_depth() function. Beta diversity was assessed using Bray-Curtis distances calculated upon these data as inputs. A Principal Coordinates Analysis (PCoA) biplot was created using data from phyloseq’s ordinate() function to examine the distribution of the samples along the first two principle coordinates. Beta dispersions of the samples were calculated using betadisper() from vegan and statistically significant pairwise differences in these were assessed using TukeyHSD() (Tukey’s Honestly Significant Differences) from the stats package. Pairwise Permutational Multivariate Analysis of Variance (PERMANOVA) between all treatments was conducted using the pairwise.adonis() function from the pairwiseAdonis v0.4.1 package (Arbizu 2020), with Bray–Curtis distances as the input variable, 9999 permutations, and Benjamini–Hochberg (BH) multiple testing p-value correction. Alpha and beta diversity of the soil microbiome samples were analyzed with 1,390 and 1,246 operational taxonomic units (OTUs) across 20 samples, respectively.

### 3.10. Leaf transcriptome analysis

Sequenced reads were trimmed using Trimmomatic (Bolger et al. 2014) with the default parameters, and mapped to the barley reference genome *Hordeum vulgare* cv. Morex genome assembly v3 using HISAT2 (Kim et al. 2015). FeatureCounts (Liao et al. 2014) was used to assess the number of reads per gene.

Differential gene expression analysis was performed in R using the following pipeline. Gene counts obtained by featureCounts were used for differential expression analysis using the DESeq2 package (Love et al. 2014) and DESeqDataSet object was constructed using the metadata table containing treatment information. Differential gene expression between different treatments was determined by fitting a negative binominal generalized linear model and performing Wald tests for the pairwise comparisons. The obtained *p*-values were adjusted using the Benjamini–Hochberg procedure and genes with *p*-value < 0.05 and an absolute log_2_ fold change ≥ 1 were considered to be significantly differentially expressed. Initially, a set of the 200 genes showing the strongest variation associated with the treatment was determined by using a variance-partitioning analysis. The z-scores for all treatments were then compared, and a heatmap showing the 50 genes with the strongest differences was created using the ComplexHeatmap package (Gu et al. 2017).

### 3.11. Rhizosphere metatranscriptome analysis

The raw metatranscriptomic reads were trimmed using Trimmomatic in paired-end mode and then processed in sequential mode with SqueezeMeta v1.6.2 (Tamames and Puente-Sánchez, 2019). Contigs were assembled *de novo* using MEGAHIT (k-mer sizes 41,61,81,99,121), and short contigs (<200 bp) were filtered using prinseq. Taxonomic and functional annotations (minimum 50% identity) involved ORF prediction using Prodigal, rRNA/tRNA detection using Barrnap and Aragorn, and DIAMOND similarity searches against GenBank, eggNOG, and KEGG, supplemented by HMMER3 for Pfam. Read mapping for quantification was performed using Bowtie2, with default parameters.

The SQM files generated by the pipeline were imported into R and processed using SQMtools (Puente-Sánchez et al. 2020) and allied packages. These files contained read counts attributed to taxonomic annotations as well as functional annotations (i.e., features) against the KEGG, COG, and Pfam databases. Prior to analysis, raw counts (abundances) were converted to per-sample relative abundances (with non-finite values being set to zero) and filtered to retain features with at least 1x10^6^ relative abundance in at least 70% of the samples. Top ten functional features for the KEGG, COG, and Pfam databases grouped by treatment were visualized using the plotFunctions() function from SQMtools using TPM (transcripts per million) values as input. Phylum-level taxonomy summaries grouped by treatment were generated using plotTaxonomy() from SQMtools with the TPM values also. Bray-Curtis distances computed from the relative abundance values calculated earlier were then used to assess beta diversity and dispersion which were then visualized using a PCoA (using pcoa() from ape (Paradis et al. 2004)) and tested for statistical significance using TukeyHSD() as described earlier. A whole model PERMANOVA as well as pairwise PERMANOVAs were also conducted as described on the Bray-Curtis distances. Finally, differential gene expression analysis was performed using DESeq2 with default parameters (Love et al. 2014).

## 4 Results

### 4.1 Barley and wheat SynComs reduce propagation of *Blumeria graminis f. sp. hordei (Bgh)* in barley leaves

To investigate whether barley exhibits induced systemic resistance when interacting with the SynComs, its roots were inoculated with barley or wheat rhizosphere SynComs, or WCS417r or a corresponding mock control solution. The barley seedlings were then grown in open-top pots with no visible changes in plant phenotypes between different treatments (Figure 2A). After 21 days, the barley seedlings were inoculated with *Bgh,* the propagation of which was monitored at seven days post inoculation (dpi). Differences in *Bgh* growth were visualized by fluorescence microscopy following DAF-FM DA staining of fungal hyphae (Figure 2B). Plants inoculated with the three inoculants exhibited comparable and significantly lower fluorescence intensities than the mock-inoculated group (Figure 2C). These results show that the barley and wheat SynComs induce ISR in a manner similar to the ISR eliciting bacterium WCS417r.

**Figure 2.**
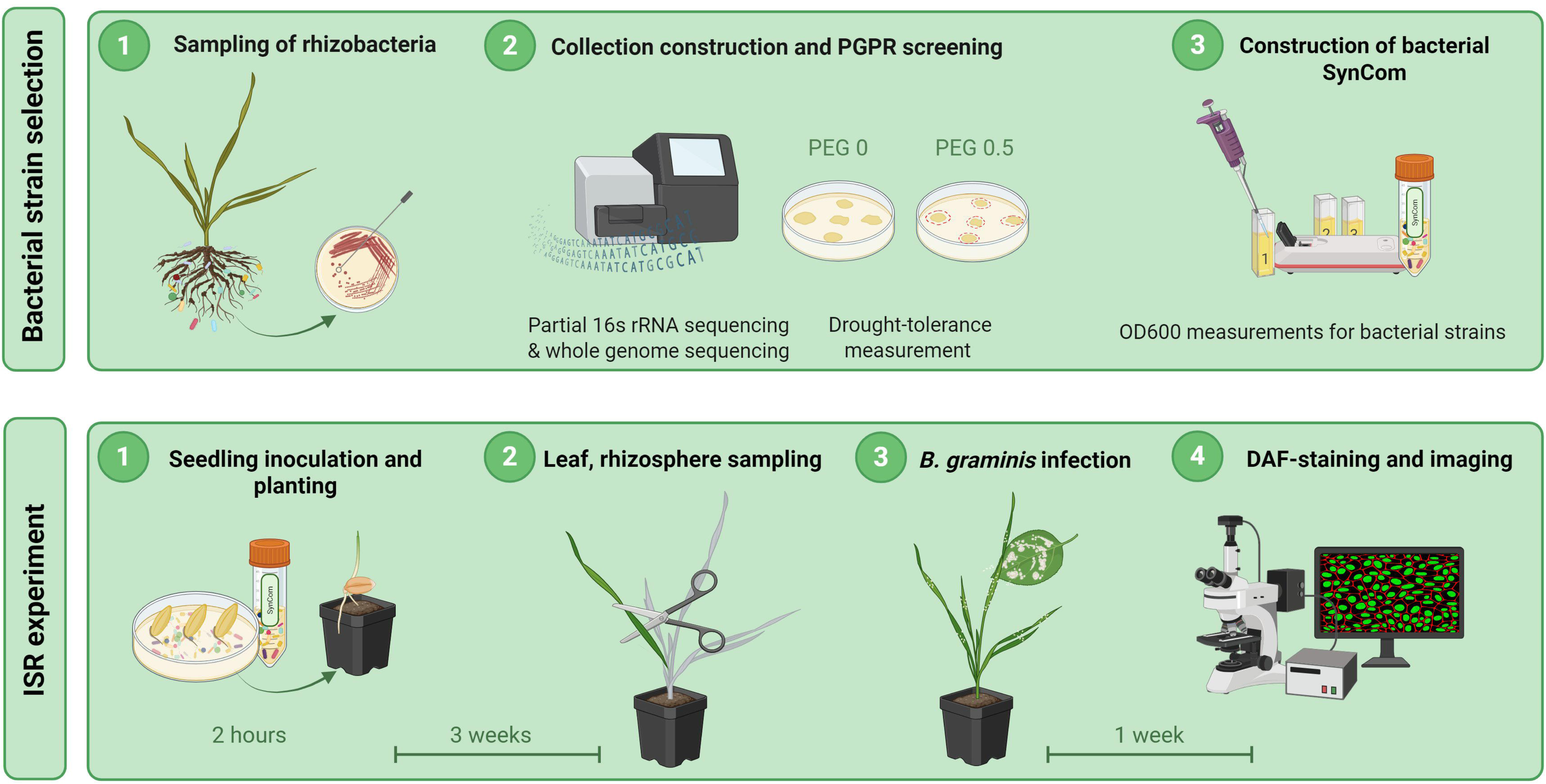
Barley and wheat SynComs enhance resistance against barley powdery mildew (*Bgh*). (A) Barley plants prior to Bgh infection, inoculated with either mock, WCS417r, wheat SynCom or barley SynCom inocula. (B) Representative fluorescence microscopy images of *Bgh* hyphae on leaf discs stained with DAF-FM-DA at 7 days post inoculation (dpi) in seedlings treated with mock, WCS417r, wheat SynCom and barley SynCom. (C) Quantification of *Bgh* on DAF-FM-DA-stained leaf discs. The *Bgh*-associated relative fluorescence units (RFU) were calculated by normalizing the measured fluorescence values against those of the uninfected controls (relative fluorescence intensity; RFU_infected / RFU_uninfected) and are shown as log₁₀(RFU_infected / RFU_uninfected). Each data point represents a single measurement from a single experiment. The experiment was repeated five times with comparable results. Different letters indicate significant differences (*p* < 0.05) according to the ANOVA and Tukey HSD tests.

### 4.2 Microbial community composition in barley rhizospheres

Next, we investigated whether increased tolerance to powdery mildew is mediated by changes in the composition of the rhizosphere bacterial community, using amplicon sequencing of the partial 16S rDNA gene. The rhizosphere microbiome comprised 1390 ASVs, which were dominated by Proteobacteria, Actinobacteriota, Bacteroidota, Firmicutes, Planctomycetota, Chloroflexi, Verrucomicrobiota, Myxococcota and Acidobacteriota, respectively (Supplementary Figure S1A). There was no difference in either alpha diversity (see Supplementary Figure S1B) or beta diversity between treatments and the control, as supported by PCoA ordination plot (Supplementary Figure S1C) and pairwise comparisons of beta dispersion (Supplementary Figure S1D Table S3). Identical ASVs to the partial 16S rRNA gene sequences of the inoculated strains were detected in the rhizosphere microbiomes (see Table S4, sheet ‘taxtable’, column ‘insyncom’), indicating that the 16S rDNA identities were present at harvest.

### 4.3 Barley leaf transcriptomes show no major response to inoculation (PERMANOVA)

Plant transcriptome sequencing was carried out to investigate barley leaf gene expression when inoculated with either MgCl_2_ (10 mM) (mock control solution), WCS417r or SynCom consisting of strains from either barley or wheat rhizosphere. Multivariate analysis of gene expression profiles after batch correction using PERMANOVA did not reveal significant differences between treatments (adjusted *p* = 0.101–0.235, R² = 0.129–0.198), indicating that treatment explained only a small proportion of the variation (Supplementary Figure S2, Supplementary Table S5).

### 4.4 Limited contribution of treatment to variance in barley leaf gene expression

To look at the sources of variability in the transcriptomic data, we quantified the variance explained by the treatment, sampling date, and residuals. Due to a strong batch effect association with sampling date, variance partitioning was interpreted after batch effect correction (Figure 3A). The batch effect correction for the sampling date reduced its caused variance to near zero, confirming the batch effect caused by the sampling date. While the treatment effect increased slightly, with a broader distribution of explained variance (typically around 12% but extending up to 60% for a subset of genes) and the residual variance decreased accordingly, the residual variance was still the main cause of variability between different treatments, despite the explanatory power of treatment increasing after the batch effect correction.

**Figure 3.**
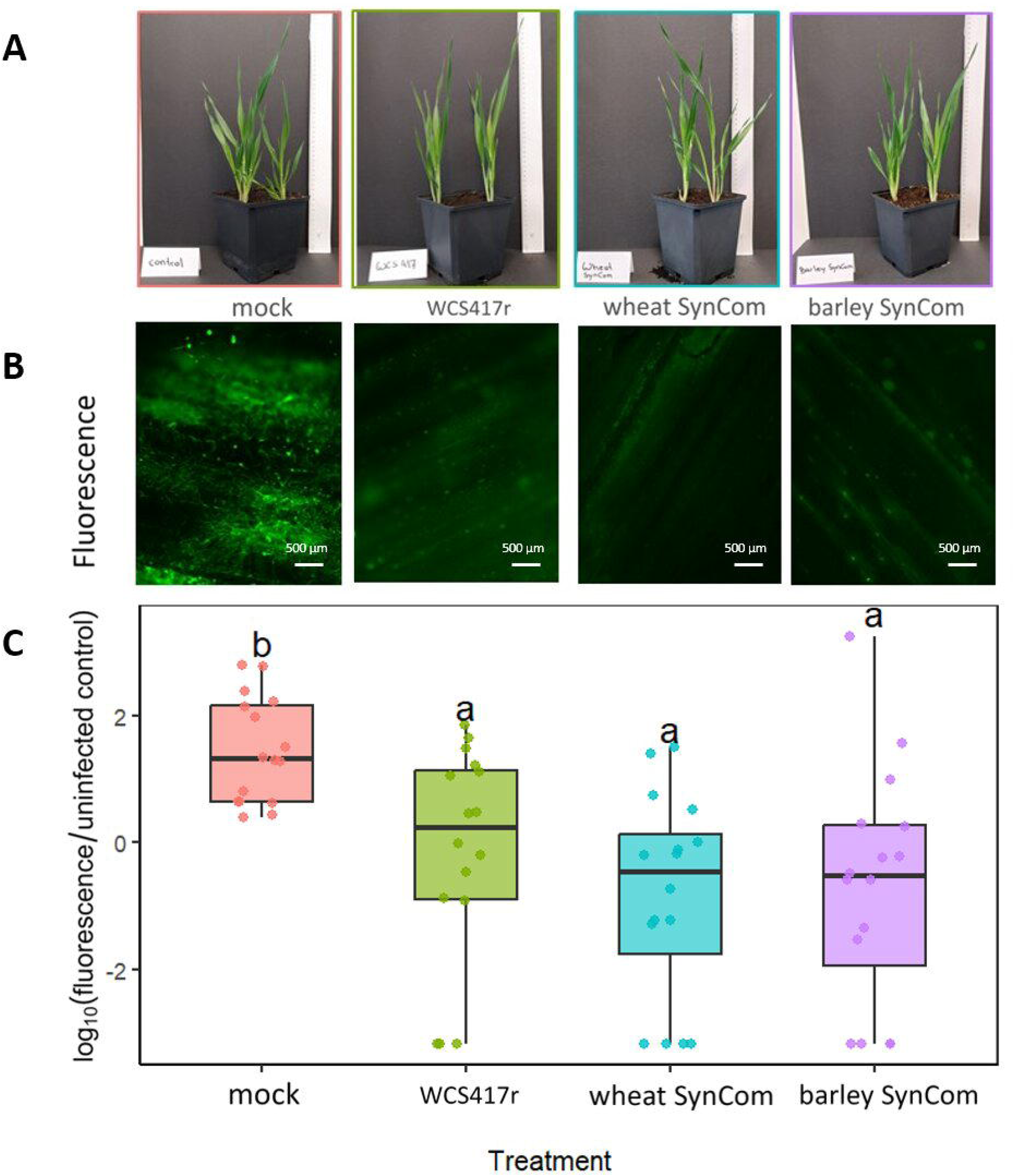
Variance partitioning of gene expression by treatment in barley leaves subjected to four treatments (mock/control, WCS417r, barley SynCom, wheat SynCom). (A) Violin plot showing variance partitioning in barley leaf transcriptomes. The violin plots show the distribution of variance explained across all genes by (i) treatment and (ii) residuals (unexplained variance). Wider regions indicate a greater density of genes with the corresponding proportion of variance explained. Treatments accounted for a small proportion of variance, while most gene expression variation remained unexplained (residuals). (B) Bar plot showing the top ten genes with the highest variance explained by treatment. The x-axis shows the percentage of variance explained, the y-axis lists the gene IDs. Each bar is divided into the variance explained by treatment (pink), date (blue) and by residuals (grey), highlighting how strongly each gene’s expression is associated with treatment. (C) VST-normalised expression of the four most treatment-associated genes. Boxplots show expression values (x-axis) across the four treatments, each panel represents one gene, illustrating differences in expression patterns between treatments.

The top 10 genes for which differences in gene expression were most strongly explained by the treatment had variance explained values of around 78% and 90% (Figure 3B). The first four of these genes (*HORVU.MOREX.r3.7HG0734910, HORVU.MOREX.r3.4HG0344820, HORVU.MOREX.r3.3HG0248440, HORVU.MOREX.r3.7HG0659040*) exhibited different expression patterns between treatments (Figure 3C). These results suggest that, after correcting for the batch effect, treatment effects on the expression of the top four genes were relatively small.

### 4.5 Differential gene expression analysis of barley leaves under different PGB treatments

To investigate transcriptional changes in barley leaves caused by the inoculations, we performed differential expression analysis. Inoculation with Barley SynCom, Wheat SynCom and WCS417r, resulted in statistically significant sets of differentially expressed genes (7, 54, and 110, respectively), however, these changes were small in magnitude related to the whole transcriptome level. Permutational analysis of variance (PERMANOVA) showed that treatment did not have a significant effect on barley leaf gene expression (adjusted *p* = 0.69–0.94; Supplementary Table S3). The pairwise comparisons showed low explanatory power (R² = 0.015–0.023), confirming that treatment accounted for only a small fraction of the total variance in gene expression. DESeq2 analysis showed that the fewest differentially expressed genes (DEGs) (7 genes) were found in the comparison between barley SynCom and the control treatment. This was followed by the comparison between wheat SynCom and the control (54 genes), and WCS417r and the control (110 genes) (Supplementary Tables S6, S7, S8). Variance partitioning analysis was used to further explore these subtle transcriptional responses. Figure 4 summarizes the 50 genes showing the strongest treatment-associated expression response in this dataset, while overall effects remained small at the transcriptome level.

**Figure 4.**
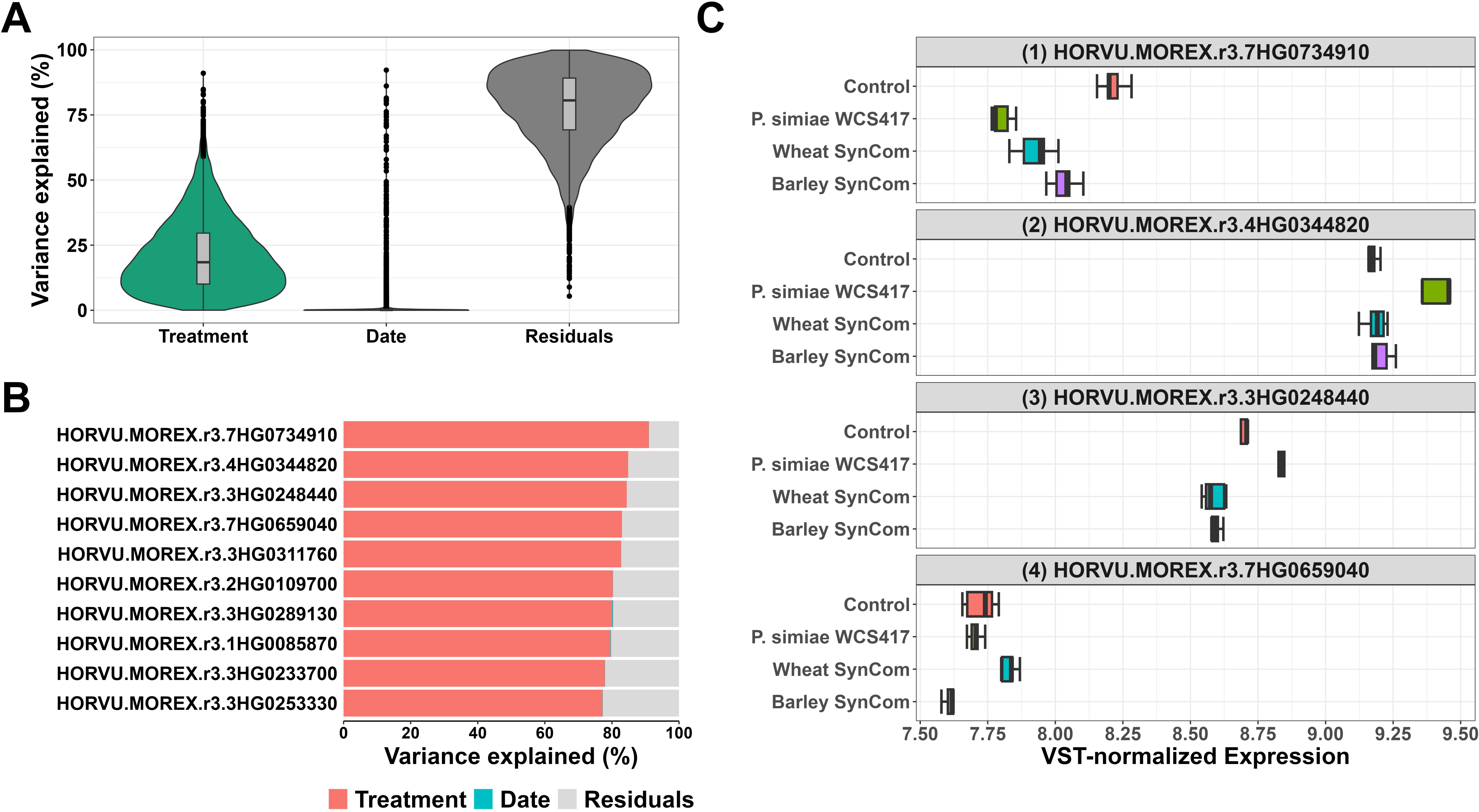
**A clustered heatmap showing normalized gene expression levels** of 50 out of 200 genes that explain the maximum variance in barley leaf gene expression after inoculation with WCS417r, barley SynCom or wheat SynCom.

The selected genes were assigned to functional categories based on their annotations and the most represented categories included Metabolism & Energy (10 genes), Transcriptional & Chromatin Regulation (8 genes), and Transport & Membrane Processes (7 genes). These categories include genes related to primary metabolism, regulatory processes, and transport functions and are often linked to plant stress adaptation and signaling. In barley SynCom inoculated plants, genes encoding a FAR1-related sequence 3, SCAR family protein, RNA-binding protein, and a translation elongation/initiation factor showed higher relative expression, while genes encoding a MYB and a Dof zinc finger protein showed lower relative expression. In contrast, WCS417r inoculation induced higher expression of mitochondrial carrier family and LanC-like protein genes, whereas a metalloprotease FtsH gene was relatively downregulated. The treatment with wheat SynCom resulted in higher expression of ERAD E3 ubiquitin ligase HRD3A and E3 ubiquitin-protein ligase genes, both involved in protein turnover and stress response. To provide an overview of the potential biological functions of these DEGs, a summary of the corresponding GO term enrichment is presented in Supplementary Figure S3.

WCS417r triggered transcriptional changes have also been described by a recent study (Sommer et al. 2026). DEG counts are sensitive to experimental and analytical approaches, therefore differences between studies are expected, hence we interpret our findings within the context of this dataset.

### 4.6 Rhizosphere metatranscriptomes show similar taxonomic and functional profiles across treatments

Metatranscriptomic analysis revealed transcripts assigned to 14 phyla across the four treatments (Control, WCS417r, Barley SynCom, Wheat SynCom). No coding sequence (CDS) was annotated for the largest proportion of reads (∼30-60%), followed by “Unclassified” and reads mapped to the plant host (Streptophyta). Bacterial phyla were detected at lower relative abundance, and most appeared distributed evenly across treatments. Overall, no consistent differences in taxonomic composition at the phylum level were observed between treatments (Figure 5A). The metatranscriptomic contigs were assigned based on sequence similarity and not reference SynCom genomes, thus taxonomy was only assigned to broad bacterial groups, and therefore read counts could not be attributed to the individual SynCom strains.

**Figure 5.**
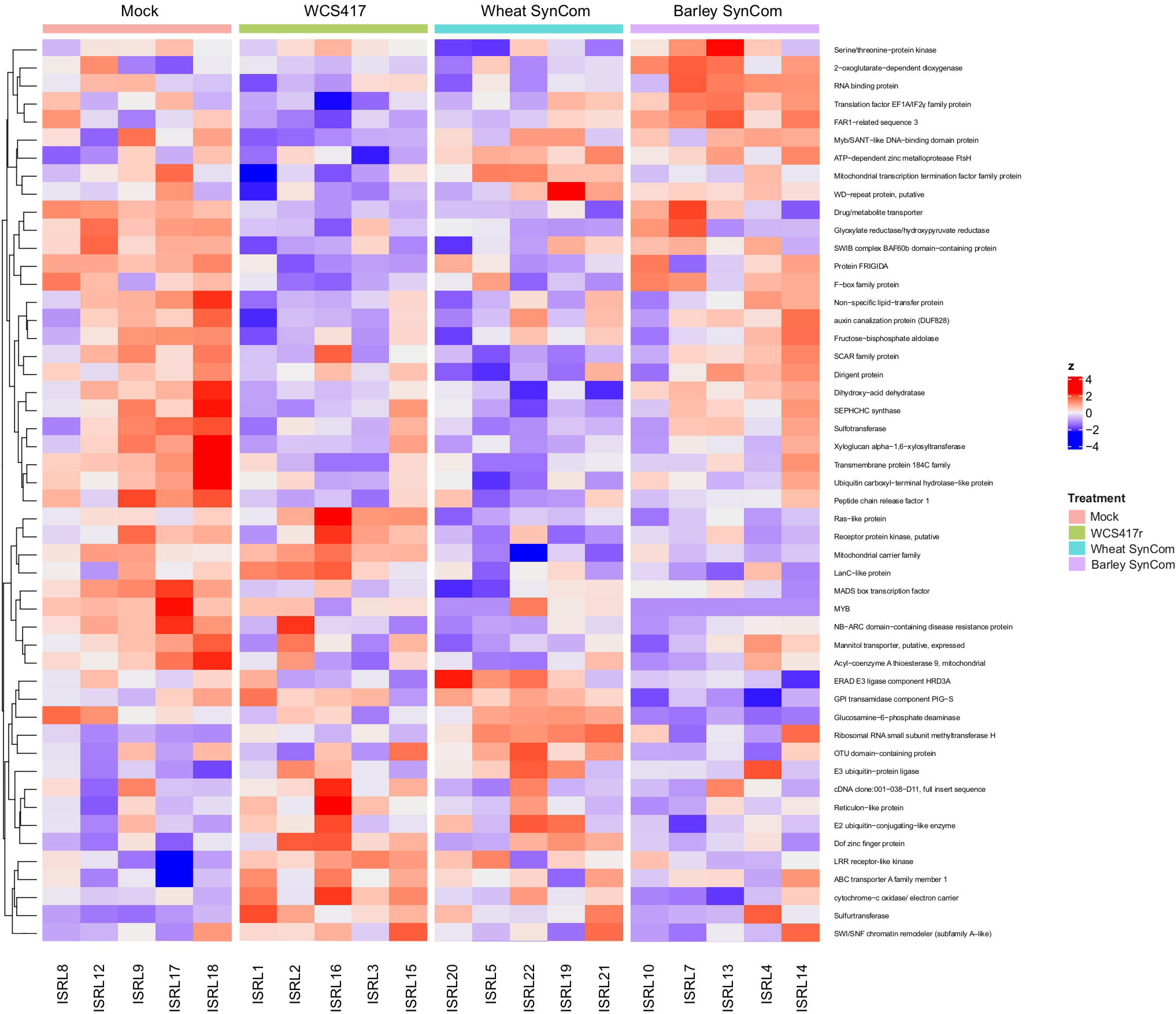
Metatranscriptomic profiles of the barley rhizosphere inoculated with WCS417r, Barley SynCom, and Wheat SynCom compared to controls. (A) Taxonomic composition across different treatments, showing the relative abundance (%) of 13 phyla in five replicates per treatment. (B) Principal coordinate analysis (PCoA) based on KEGG (A), COG (B), and Pfam (C) annotations. Axes indicate the percentage of variance explained by PC1 and PC2. (C) Alpha diversity calculated from KEGG, COG, and Pfam functional annotations, represented by Shannon index, richness, and evenness (nine plots total). The y-axis shows the corresponding diversity metric, while the x-axis shows replicates per treatment.

Principal coordinate analysis (PCoA) of KEGG (A), COG (B), and Pfam (C) annotations showed that despite few outliers from Control plants, samples from all treatments were mostly grouped together, with ellipses overlapping in all three plots (Figure 5B). The explained variance was similar between annotations, with the highest for KEGG (PC1: 48.5%, PC2: 20.2%) and lower for COG (PC1: 43.4% and PC2: 21.%) and Pfam (PC1: 55%, PC2: 7.2%).

Alpha diversity was measured with Shannon index, richness, and evenness using KEGG, COG, and Pfam annotations (Figure 5C). The diversity values were similar between all treatments, and statistical tests showed no significant differences between all pairwise comparisons (abbreviated ns). Few outliers and variation was present, particularly for Wheat SynCom and Control samples, but no clear pattern related to treatment was detected.

### 4.7 Differentially expressed functional features in rhizosphere metatranscriptomes

Differential gene expression analysis of the metatranscriptomes revealed 14 functional features (KEGG, COG, and PFAM terms) to be enriched across the bioinoculant applications compared to the controls (BH adjusted *p*-value <= 0.05 & absolute log_2_-fold change >= 1) (Figure 6A). Application of WCS417r was associated with upregulation of genes annotated with the following features: K03520 (KEGG), COG1529 (COG), PF01778 (PFAM), and PF20256 (PFAM). These annotations include proteins associated with redox-related metabolic functions, ribosomal components, and molybdenum cofactor-binding domains, In addition, K01919 (glutamate-cysteine ligase) and K05516 (stress-induced DnaJ-like molecular chaperone) were also upregulated under WCS417r treatment.

**Figure 6.**
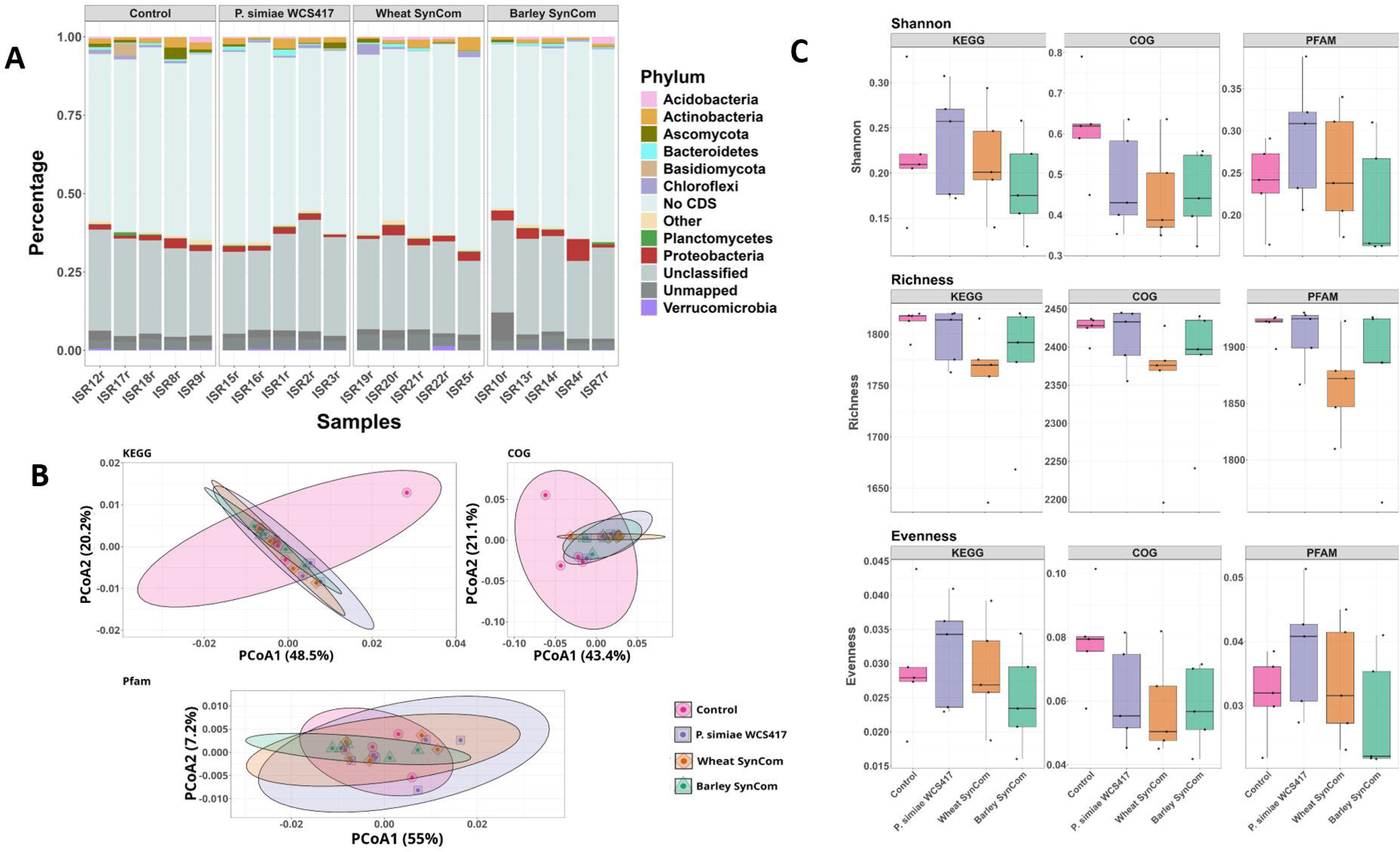
Differential gene expression analysis of barley rhizosphere metatranscriptomes highlighting significantly upregulated and downregulated functional features. (A) Volcano plots showing significantly upregulated and downregulated terms across KEGG, COG, and PFAM categories. The x-axis represents log₂(fold change), and the y-axis represents −log₁₀(BH-adjusted *p* value). Significantly changed terms are highlighted compared to all other terms within each category. (B) Boxplots of representative significantly upregulated and downregulated terms, displayed by treatment condition. Each boxplot shows log₁₀(TPM) values for the Control versus the respective treatment (e.g., KEGG term K05516 in Control vs. Barley SynCom).

In contrast, several metatranscriptomic features were downregulated under WCS417r treatment, including ENOG4112351 (eggNOG), ENOG410ZRFR (eggNOG), and PF02868 (PFAM). ENOG4112351 and ENOG410ZRFR are annotated as proteins of unknown or uncharacterized function, while PF02868 corresponds to a thermolysin metallopeptidase–related domain.

Application of the wheat SynCom resulted in statistically significant downregulation of PF02868; no KEGG or eggNOG features were significantly downregulated in this treatment. In contrast, K05516 (KEGG; stress-induced DnaJ-like molecular chaperone) and PF01614 (PFAM; IclR-family transcriptional regulator, C-terminal domain) were significantly upregulated under wheat SynCom application.

Application of the barley SynCom was associated with the upregulation of genes annotated with K02109 (KEGG; F-type H⁺-transporting ATPase subunit b), COG0711 (COG; ATP synthase–related protein), and PF01061 (PFAM; ABC-2 type transporter). In addition, K05516 (KEGG; stress-induced DnaJ-like molecular chaperone) and ENOG410XQKZ (eggNOG; oxidoreductase-related domain protein) were significantly upregulated under barley SynCom application.

Several features were downregulated in the barley SynCom treatment, including ENOG4112351 (eggNOG; protein of unknown function) and PF02868 (PFAM; thermolysin metallopeptidase–related domain).

Comparison across treatments revealed that PF02868 was significantly downregulated and K05516 significantly upregulated in all three treatments (WCS417r, barley SynCom, and wheat SynCom). ENOG4112351 was downregulated under WCS417r and barley SynCom application but not under the wheat SynCom treatment.

## 5 Discussion

In this study, we investigated the ability of SynComs, which are composed of native bacteria from barley and wheat rhizospheres, to promote resistancein barley plants against barley pathogen *Blumeria graminis* f. sp. *hordei (Bgh)*. SynCom-trigerred ISR reduced *Bgh* propagation in infected plant leaves, which is consistent with the positive effect of the model strain WCS417r, which is known to induce ISR in barley (Sommer et al. 2026) and is therefore used as a positive control in ISR experiments.

### 5.1 The potential of SynCom as a biological control agent against plant pathogens

SynComs are used in studies of plant-microbe interactions and have the potential to be used as bioinoculants, as they can enhance crop disease resistance and resilience under environmental stress, reduce the need for chemical pesticides and modulate host phenotype (Naik et al. 2019; Vorholt et al. 2017; Chesneau et al. 2025). Unlike single-strain inoculants, SynComs more accurately reflect the complexity of a natural microbiome and have sometimes been shown to have a stronger effect on plant health than individual strains applied alone (Liu et al. 2023). SynComs are often assembled using rhizobacteria, whose composition differs from the bulk soil microbiome, partly due to root exudates released by the plant (Shayanthan et al. 2022; Koprivova and Kopriva 2022). A certain proportion of these strains are root-associated plant growth promoting rhizobacteria. Among other plant beneficial traits, some of them activate plant immune responses through ISR, a mechanism triggered in the plant root that can protect above-ground parts of the plant rom pathogen invasion (Pieterse et al. 2014).

The potential to act as a biocontrol agent against plant pathogens has been well studied using the model strain WCS417r (Pieterse et al. 2021; Sommer et al. 2024; Vorholt et al. 2017). In addition, there are studies applying SynComs for crop resistance. For instance, the application of a SynCom consisting of bacterial isolates from the endosphere of the maize variety Jala landrace was found to improve plant growth and inhibit the growth of phytopathogenic fungi, including *Pestalotia* sp. and *Colletotrichum* sp. (La Vega-Camarillo et al. 2023).

Furthermore, it has been reported that microbial inoculants can help plants tolerate abiotic stress. This is achieved by increasing nutrient availability, fixing nitrogen, producing plant growth promoting (PGP) hormones such as indole-3-acetic acid, and decreasing ethylene production (Mastouri et al. 2010; Kejela 2023; Pan and Cai 2023; Aasfar et al. 2024). These additional characteristics can promote plant growth in challenging environmental conditions when employed in SynComs (Wallenstein 2017; Shayanthan et al. 2022; Rigerte et al. 2025). The barley SynCom used here was previously characterized in the context of drought tolerance (Rigerte et al. 2025). The relationship between drought tolerance and ISR is still to be explored, however this broader potential further strengthens the idea of SynComs as a sustainable tool in agriculture, not only against pathogens but also under future climate stress scenarios.

Studies using SynComs as bioinoculants are thus far limited, and their outcomes often depend on the host plant, the pathogen and environmental conditions. This is demonstrated by the context-dependent effects of SynComs on disease progression and microbiome composition (Pfeilmeier et al. 2024; Liu et al. 2023). Transferring successful results from *in vitro* or greenhouse experiments to the field is challenging due to the interaction with the native microbiome and other factors that can strongly influence SynCom performance. Still, SynCom approaches are a promising way forward for sustainable alternatives to chemical pesticides. Our work adds to this evidence by testing their potential in the barley–*Bgh* system. Our data reveal that exposure of barley roots to either barley- or wheat-derived SynComs enhanced the resistance of the above-ground tissues of the plants against powdery mildew, suggesting the induction of ISR by the SynComs used.

### 5.2 Reduced fungal growth in barley leaves after SynCom and WCS417r treatments

Our DAF staining results show that inoculation with both SynComs and WCS417r reduced fungal growth in barley leaves following infection with *Bgh*, showing no major difference between different inoculants. Lower fluorescence levels observed in our treatments suggest that plants inoculated with beneficial bacteria had reduced pathogen growth (Lenk et al. 2018; Lenk et al. 2019; Brambilla et al. 2022; Brambilla et al. 2023).

Restriction of fungal growth is a common outcome of ISR, in which root-associated beneficial bacteria prime the plant for enhanced defense response upon pathogen attack which results in reduced colonization instead of complete resistance (van der Ent et al. 2009; Pieterse et al. 2014). In barley, reduced hyphal growth and colony establishment of *Bgh* are well-established phenotypic indicators of enhanced resistance and are widely used to quantify disease suppression in both host and non-host interaction studies (Hückelhoven et al. 2001; Panstruga and Schulze-Lefert 2002).

The reduced fungal growth after inoculation with SynCom and WCS417r suggests that these treatments enhance plant’s ability to restrict pathogen development in leaf tissues. Such results are consistent with ISR-mediated priming in which immune responses are deployed more rapidly upon pathogen attack, resulting in lower pathogen biomass (Pieterse et al. 2014). Together, these results suggest that both SynComs and the model strain WCS417r limit the growth of *Bgh* in barley leaves, highlighting their potential usage in microbiome-mediated disease suppression.

### 5.3 Limited effects of synCom inoculation on barley rhizosphere microbiota

The alpha or beta diversity of the barley rhizosphere prokaryotes did change significantly between treatments, suggesting that the addition of SynCom did not significantly alter richness or community composition. The relative abundances of bacterial ASVs in rhizosphere samples remained consistent across batches of the same treatment. The available literature on changes to the microbiome after inoculation with plant beneficial bacteria (PBB) is inconclusive. For example, (Hao et al. 2023) demonstrated adding a SynCom to desert soil containing a mixture of tall fescue and ryegrass resulted in an increase in beneficial bacteria and greater bacterial diversity. In contrast, (Qiao et al. 2024) reported that SynCom addition to grafted watermelon plants led to a significant increase in the relative abundance of *Pseudomonas* in the rhizosphere. Conversely, research on *Zea mays* B73 revealed that the seed inoculation with PGP bacterium mitigated drought stress in maize without altering the composition of the rhizosphere bacterial community (Yim et al. 2025). Similarly, (Montoya et al. 2025) demonstrated that although inoculation of tomato seedlings with a bacterial SynCom induced the expression of numerous genes in the roots and shoots, the composition of the rhizosphere communities was not significantly altered.

### 5.4 Subtle but specific transcriptional responses in barley leaves Following SynCom and WCS417r inoculation

Minimal transcriptomic differences in barley leaves were observed for all four inoculations. Treatment explained only a minor proportion of the variance for most genes, and PERMANOVA or GO enrichment analyses didn’t detected clear separation between Control, WCS417r and both SynComs. This suggests that most of the systemic transcriptional responses in leaves are subtle, affecting only a small number of genes. This is consistent with the recent findings of A. S. and A.C.V. (Sommer et al. 2026) and the existing literature. For example, none of the genes tested in *A. thaliana* leaves showed a consistent change in gene expression during WCS417r-mediated ISR (Verhagen et al. 2004), suggesting that ISR activation in plant leaves is not associated with strong changes in gene expression. Similarly, a study on the barley root colonization of a beneficial endophyte *P. indica* showed clear physiological evidence of enhanced systemic resistance against *Bgh*, without resulting in strong transcriptional response in barley leaves under non-challenged conditions (Molitor et al. 2011). Furthermore, barley growth-promoting and fungal-root-pathogen-inhibiting effects of *Sebacina vermifera* and rhizosphere bacteria are accompanied by a mild transcriptional response in the host upon early colonization by *S. vermifera* and bacteria (Mahdi et al. 2022). The findings from literature and our results are consistent with the priming effect associated with ISR (Conrath et al. 2001; Conrath et al. 2015; Sommer et al. 2026).

Our analysis of differentially expressed genes showed that mock treated plants exhibited higher activation of several defense-related genes, including those encoding MYB, NB-ARC proteins, PPR proteins, as well as lipid- and cell wall–related enzymes. As these samples were collected prior to pathogen inoculation, this elevated defense gene expression may reflect a higher basal stress or defense state in mock inoculated plants. In comparison, SynCom inoculation reduced the expression of these genes, which mat reflect an ISR-like priming, where the plants show lower basal defense gene expression while remaining prepared for a response under pathogen infection. WCS417r and both SynComs also influenced several metabolic and signaling pathways, for example, Barley and Wheat SynCom both downregulated phosphatidate cytidylyltransferase and phosphoglucose isomerase 1, indicating changes in lipid and central carbon metabolism during microbe-mediated priming. These enzymes typically increase during strong plant defense activation, suggesting that SynCom inoculated plants invested reduced amount of lipid and carbon resources into plant basal defense. In addition, WCS417r upregulated LEA proteins and MTA cycle–related enzymes, while both WCS417r and Barley SynCom increased laccase and MYB-like transcription factor expression, which may support cell-wall–related defense. Together, these results suggestthat different beneficial bacteria activate partially overlapping but distinct transcriptional responses linked to ISR and pathogen protection.

Our leaf transcriptome results are in line with the literature. For example, (van de Mortel et al. 2012) found that *A. thaliana* root inoculation with WCS417r did not result in consistent transcriptional changes in plant leaves prior to the infection, which suggests that ISR results via priming rather than changes in plant gene expression. Similarly, (Lakkis et al. 2019) studied the ability of *Pseudomonas fluorescens* PTA-CT2 to induce ISR in grapevine plants against *P. viticola* and *B. cinerea* and demonstrated that plant inoculation did not result in consistent changes of basal defenses prior the infection. To conclude, these results highlight the importance of sampling during pathogen infection to further investigate how PGP treatments modulate ISR against *Bgh*.

### 5.5 Rhizosphere microbiome gene expression analysis detects treatment-specific and shared functional changes

Metatranscriptomics offers a practical way of discovering the mechanisms underlying the ISR process, rather than merely identifying genes in bacterial genomes that may contribute to it (Yang et al. 2025). In all three treatments, two features were consistently affected: K05516 (stress-induced DnaJ-like molecular chaperone), which was upregulated, and PF02868 (thermolysin metallopeptidase–related domain), which was downregulated. This impliesthat the application of these bioinoculants elicited a shared microbial response involving stress adaptation and changes in specific functional activities (Yamashino et al. 1994). The wheat SynCom treatment induced fewer transcriptional changes in barley rhizosphere compared to the other treatments, but included the upregulation of PF01614, a bacterial transcriptional regulator. This indicates that although fewer features were enriched, the SynCom also stimulated microbial signaling and stress response functions.

Inoculation with the barley SynCom was associated with increased expression of features annotated as ATP synthase-related proteins (K02109, COG0711) and ABC-type transporters (PF01061). This implies that inoculation with the barley SynCom may influence microbial energy related functions and transport processes in the rhizosphere. Furthermore, the upregulation of K05516 and ENOG410XQKZ suggests enhanced microbial stress adaption and redox-related functional responses under this treatment (Yamashino et al. 1994). At the same time, downregulation of PF02868 and ENOG4112351 was observed, similar to WCS417r, indicating shared regulation of specific, non-central metabolic features.

Inoculation with WCS417r was associated with the upregulation of stress related and transport associated functions, suggesting that microbial activity is functionally adjusted instead of generally increased (Andrade et al. 2023). The enrichment of detoxification and transport related functions indicates an increased microbial capacity to manage environmental stress and to support nutrient exchange, which may contribute to plant stress protection prior to pathogen infection (Andrade et al. 2023; Desrut et al. 2020). Additionally, increased expression of functions related to nitrogen metabolism and microbial signalling suggests that WCS417r promotes a more responsive and metabolically active rhizosphere microbiome (Pieterse et al. 2021).

Conversely, genes related to energy and ribosomes were downregulated in this treatment. This indicates reduced metabolic activity, likely due to the community allocating resources towards specialised functions rather than growth (Blazewicz et al. 2013).

The taxonomic overview of the metatranscriptomes showed that most of the transcripts could not be assigned to coding sequences or were put in the group “unclassified.” This is similar to earlier studies, where functional metatranscriptomic datasets often had many unannotated reads. For example, (Howe et al. 2023) described a large community of unclassified active microbes in metatranscriptomic profiles and (Jiang et al. 2016) showed that only 19–76% of mRNA reads were annotatable across diverse microbiome samples. Host-derived Streptophyta transcripts were detected and removed prior to downstream taxonomic analyses.

The observed differential gene expression patterns in the rhizosphere microbiome suggest that microbial responses to biostimulant inoculation are selective and function specific, rather than reflecting widespread changes across the entire community. For instance, a study on (RNA) of the rhizosphere microbiome of sugar beet seedlings grown in pathogen-suppressive soil showed restricted pathogen triggerred microbial response, mainly in stress- and signaling-related transcripts, further supporting the idea that inoculated treatments are more associated with specific transcripts, rather than major changes in entire microbial community (Chapelle et al. 2016; Schlatter et al. 2017). Similarly, a review on soil microbial ecology and metatranscriptomics concluded that metatranscriptomic responses are most apparent under strong environmental changes, while weaker disturbances result in milder changes (Peng et al. 2024). Together these results show that inoculation with different SynComs and WCS417r strain elicits shifts in rhizosphere microbial functional activity, pointing towards modulation of energy metabolism, signalling and nutrient exchange instead of mere growth-boosting.

## Conclusions

Our study demonstrates that SynComs derived from drought stressed barley and wheat rhizospheres are capable of inducing systemic resistance in barley against powdery mildew infection. Furthermore, the wheat SynCom induced ISR response in barley that was comparable to the one induced by the barley SynCom, suggesting that ISR can be elicited by synthetic microbial communities originating from a related, but distinct host species.

Although the two SynComs were composed of strains from different genera, all bacterial members belong to PGB, e.g. *Pseudomonas, Bacillus/Priestia, Rhizobium*-related taxa, and *Streptomyces*. The overlap of the PGP traits may explain the ability of SynComs to induce ISR in barley. At the same time, treatment-specific differences observed in rhizosphere metatranscriptomes indicate that distinct microbial communities may achieve similar ISR outcomes through different functional and metabolic strategies.

A further conclusion of this study is the detection of treatment-specific shifts in rhizosphere microbial gene expression, including for the well-established ISR model strain *Pseudomonas simiae* WCS417r. These results provide novel insights into how ISR-eliciting bacteria and communities adjust rhizosphere microbial activity beyond effects on host gene expression.

Finally, application of both SynComs did not affect host gene expression, as demonstrated by the absence of differential gene expression in host leaf tissues under SynCom application. In the context of ISR, this suggests that SynCom application alters the behavior of the soil microbial community, priming the host plant to defend itself against potential pathogen attacks.

## 6 Data Availability Statement

Metatranscriptomic and 16S rRNA gene amplicon sequencing data of soil bacterial communities, as well as host leaf RNA sequencing data of barley exposed to beneficial bioinoculants, are available under NCBI BioProject ID PRJNA1405459.

## 8 Author Contributions

LR: Formal Analysis, Investigation, Methodology, Visualization, Writing – original draft, Writing – review & editing. AS: Investigation, Methodology, Writing – review & editing. ACV: Investigation, Methodology, Writing – review & editing. LDPS: Formal Analysis, Writing – review & editing. AH: Conceptualization, Funding acquisition, Writing – review & editing. TR: Conceptualization, Funding acquisition, Writing – review & editing. MTT: Conceptualization, Funding acquisition, Methodology, Project administration, Resources, Supervision, Validation, Writing – review & editing.

## 10. Acknowledgments

The authors would like to thank Alga Zuccaro for leading the DECRyPT consortium and support of our work. They would also like to thank the EVE administration at the UFZ/iDiv High-Performance Computing (HPC) cluster for maintaining the EVE Cluster, on which the bioinformatics analyses were performed. The authors would also like to express their gratitude to Kerstin Hommel and Carina Heuschmann for their invaluable assistance in the laboratory.

