## Supplementary Material for "The induction of systemic resistance to barley powdery mildew by rhizosphere bacterial communities does not disrupt the structure or function of native microbial communities"

**Supplementary Figures and Tables**

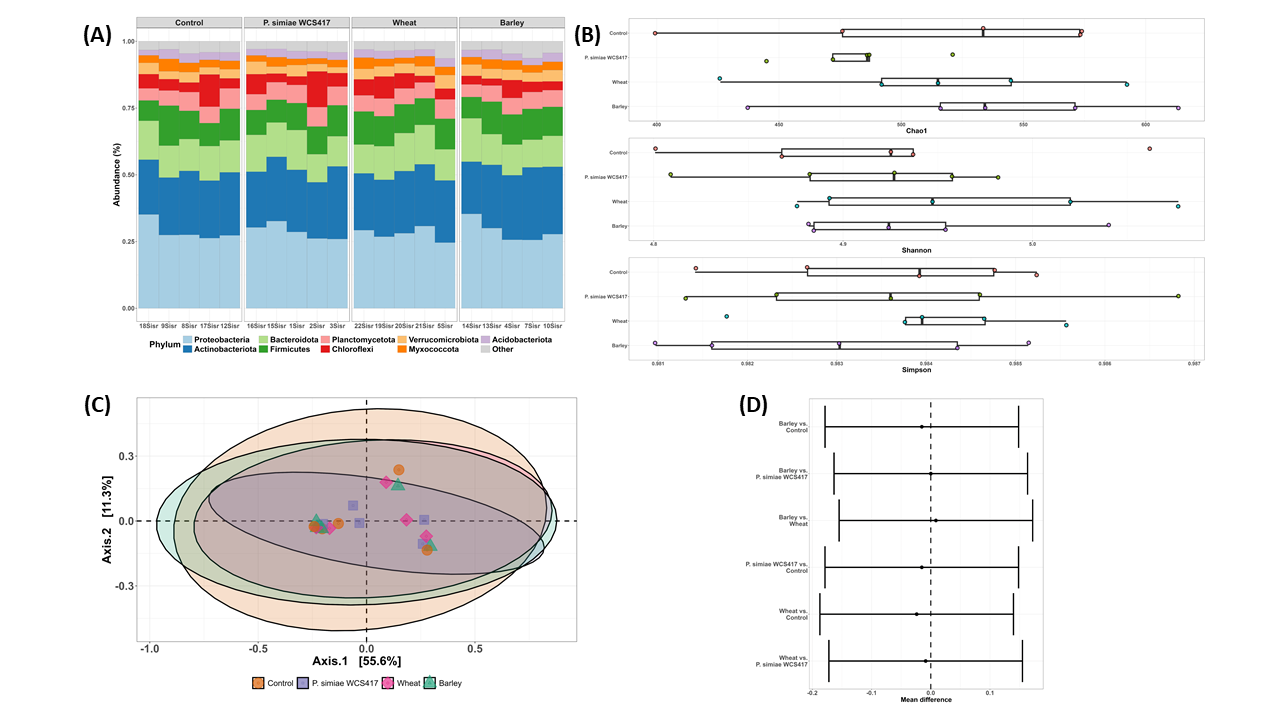

**Supplementary Figure 1. 16S rDNA amplicon sequencing results from soils of barley hosts** treated with a control treatment and three different bioinoculants (*Pseudomonas simiae* WCS417, wheat SynCom, barley SynCom) respectively. (A) Relative abundances of top 10 phyla. (B) Chao1, Shannon, and Simpson indices of alpha diversity. Statistically significant differences between treatments are marked with asterisks. (C) PCoA ordination plot of the samples colored by treatment along the first two principal coordinates. (D) Tukey’s HSD plot for all pairwise comparisons of beta dispersions in the treatments. Error bars that do not include zero (vertical dotted line) and colored red indicate statistically significant differences.

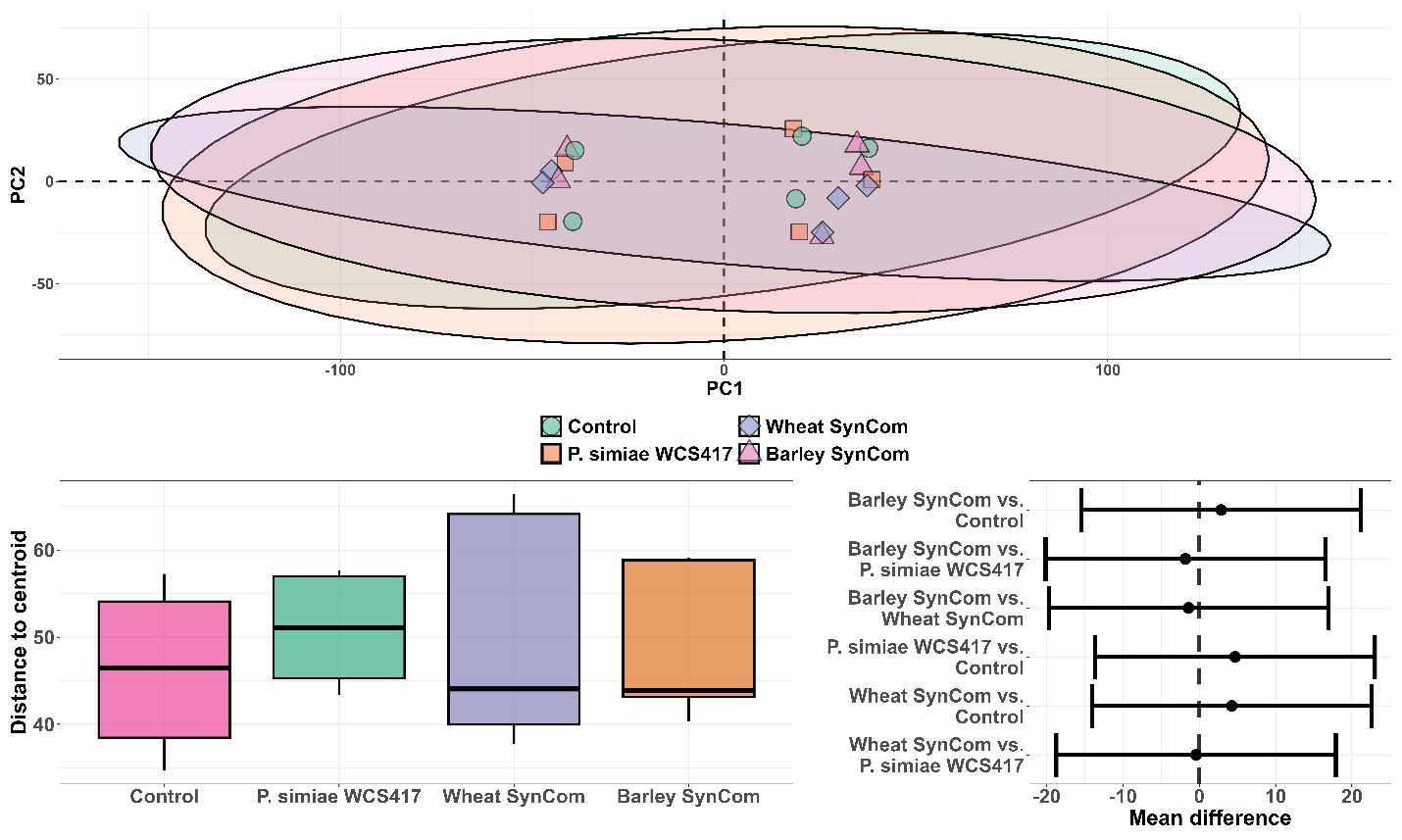

**Supplementary Figure S2.** Multivariate analysis of barley leaf transcriptomes with different inoculations (Control, WCS417, Barley SynCom, Wheat SynCom). (A) PCoA ordination plot of the samples colored by treatment along the first two principal coordinates. (B) Boxplots of the distance to centroid for each treatment, indicating no significant differences. (C) Pairwise PERMANOVA comparisons of treatments, showing mean differences close to zero.

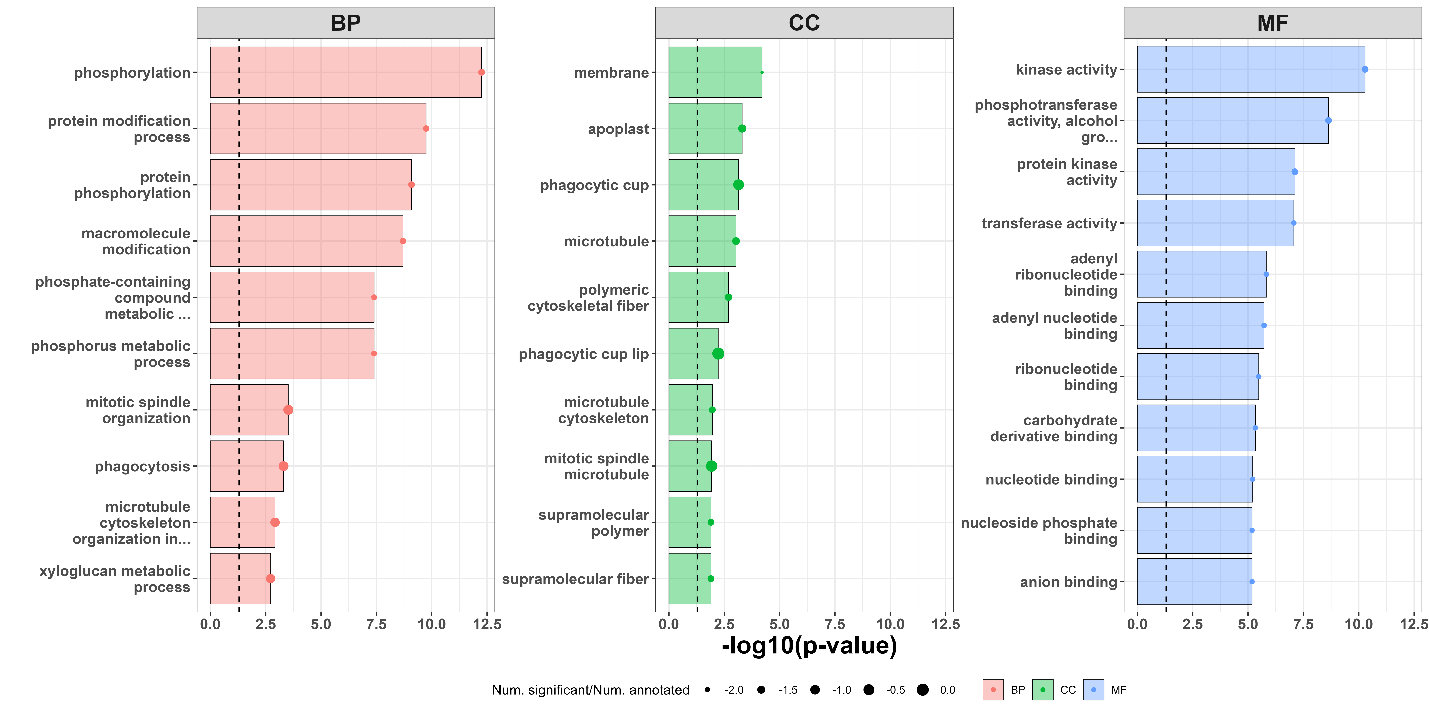

**Supplementary Figure S3.** Gene Ontology (GO) enrichment analysis of barley leaf transcriptomes inoculated with the Wheat SynCom compared to control plants. Enriched GO terms are shown for the three main categories: Biological Process, Cellular Component, and Molecular Function. The y-axis represents the GO terms, and the x-axis indicates statistical significance as –log10(p-value).
